# Replay of large-scale spatio-temporal patterns from waking during subsequent slow-wave sleep in human cortex

**DOI:** 10.1101/221275

**Authors:** Xi Jiang, Isaac Shamie, Werner Doyle, Daniel Friedman, Patricia Dugan, Orrin Devinsky, Emad Eskandar, Sydney S. Cash, Thomas Thesen, Eric Halgren

## Abstract

Animal studies support the hypothesis that in slow-wave sleep, replay of waking neocortical activity under hippocampal guidance leads to memory consolidation. However, no intracranial electrophysiological evidence for replay exists in humans. We identified consistent sequences of population firing peaks across widespread cortical regions during complete waking periods. The occurrence of these Motifs were compared between sleeps preceding the waking period (Sleep-Pre) when the Motifs were identified, and those following (Sleep-Post). In all subjects, the majority of waking Motifs (most of which were novel) had more matches in Sleep-Post than in Sleep-Pre. In rodents, hippocampal replay occurs during local sharp-wave ripples, and the associated neocortical replay tends to occur during local sleep spindles and down-to-up transitions. These waves may facilitate consolidation by sequencing cell-firing and encouraging plasticity. Similarly, we found that Motifs were coupled to neocortical spindles, down-to-up transitions, theta bursts, and hippocampal sharp-wave ripples. While Motifs occurring during cognitive task performance were more likely to have more matches in subsequent sleep, our studies provide no direct demonstration that the replay of Motifs contributes to consolidation. Nonetheless, these results confirm a core prediction of the dominant neurobiological theory of human memory consolidation.

## Introduction

“Replay” of spatio-temporal neuronal activity patterns from waking during slow-wave sleep (SWS) was first demonstrated in rat hippocampal place cells ^1,2^. Hippocampal replay is hypothesized to play a crucial role in memory consolidation, orchestrating cortical replay during sleep, resulting in synaptic plasticity and thus memory consolidation ^1,3^ Indeed, replay coordinated with the hippocampus occurs in visual ^4^ and prefrontal ^5,6^ cortices, and memory strength can be modulated by disrupting or augmenting replay ^7^. While most replay studies were conducted in rats, one study found replay of firing-patterns across motor, somatosensory, and parietal cortices in macaques ^8^, where both within-area and across-area (parietal and motor) neural ensembles were seen to reactivate. Reactivation of local hemodynamic patterns evoked by previously studied stimuli have been demonstrated in humans ^9-13^. However, hemodynamic signals are too sluggish to detect temporal (i.e. sequential activation) patterns that are critical for memory encoding and consolidation, or to allow identification of fine temporal correlations with sleep graphoelements relevant to memory processing. Replay may also occur in scalp EEG ^14^ or MEG ^15^ recordings, but their poor spatial resolution and correlation with neuronal firing limit their interpretation. Similarly, average correlation patterns of ECoG broadband gamma power over sensory areas during NREM was reported to resemble the correlation patterns during awake movie-watching periods ^16^. While supportive of spatial pattern reactivations in the human cortex specifically during NREM, such full-night average methods were insufficient for investigating whether, like animal replay, fine temporal structures/sequences could be established for widespread cortical reactivations. Here we utilized long-term (>4 days) recordings using intracranial electrodes placed for the localization of seizure onset and eloquent cortex in subjects with focal epilepsy. Electrode contacts were either within the cortex and hippocampus (stereoelectroencephalography, SEEG), or were on the cortical surface (electrocorticography, ECoG). We measured High Gamma (HG) activity (70-190 Hz) across widespread cortical locations. HG amplitude tracks local neuronal firing ^17^, and predicts declarative memory encoding ^18^ and retrieval ^19^. Across cognitive tasks and conditions, HG amplitude is strongly correlated with hemodynamics, but has much faster (millisecond) resolution ^20^.

We simultaneously recorded Local Field Potentials (LFP) from multiple cortical sites to detect the widespread cortical modulations characteristic of SWS. LFPs are generated by spatially organized and temporally-coherent trans-membrane currents, and can be seen at the scalp as EEG and MEG ^21^. The SWS characteristics we identified include sleep spindles (∼500-2000 ms bursts of 10-17 Hz waves) and delta activity ^22,23^. Delta is comprised of rhythmic “downstates” and “upstates”, wherein the cortex alternates at ∼0.5-2 Hz between near-silence and firing similar to quiet waking ^24^. Cortical replay in rodents is associated with spindles and down-to-upstates^22,23^, and memory consolidation in humans is associated with their density in the EEG ^9,25^. In addition to spindles and delta activities, whose phasic coupling to ripples appears to enhance consolidation ^26^, there have also been reports of cortical theta oscillations coordinating sequential encoding of episodic memory ^27^ and their occurrence in NREM sleep being predictive of successful verbal memory recall ^28^. We also related possible replay events to hippocampal sharp-wave ripples, which in rodents mark the occurrence of cellular replay ^29^.

Our strategy for detecting candidate spatiotemporal patterns for replay was based on several previous observations. In humans, hemodynamic patterns specific for particular memories are distributed during encoding and retrieval ^30^. In rodents, c-Fos up-regulation is widespread across multiple cortical areas after learning ^31,32^, with a delayed wave of hippocampal c-Fos expression, peaking 18-24 hours after training regimens that produce long lasting memories ^33^. Therefore, we reasoned that distributed spatio-temporal patterns of sequential HG amplitude peaks would be found if we searched in 6-19 hours of continuous waking ECoG/SEEG. Based on their spatiotemporal similarity, we clustered these patterns into recurring “Motifs.” We hypothesized that these Motifs would be more prevalent during SWS periods following the waking period (“Sleep-Post”) than those preceding it (“Sleep-Pre”), thus indicating a “replay” of the day’s neural activities during subsequent sleeps.

## Results

For each subject, widespread cortical Motifs were identified during waking, and spatiotemporal activity patterns similar to those Motifs (and therefore considered putative *reactivations* of Motifs) were identified from the two SWS periods that comprise Sleep-Post (i.e. subsequent to the waking period) and the two SWS periods that comprise Sleep-Pre (i.e. preceding the waking period). A significant majority of those Motifs were more likely to occur more often in Sleep-Post than in Sleep-Pre. Novel Motifs, or Motifs identified during active cognition, were more likely to be preferentially re-activated during Sleep-Post. Motif recurrence was associated with increased cortical spindles and slow oscillation power, as well as broadband hippocampal activation. A list of which subjects were involved in each analysis can be found as Supplementary Table S1.

**Figure 1.**
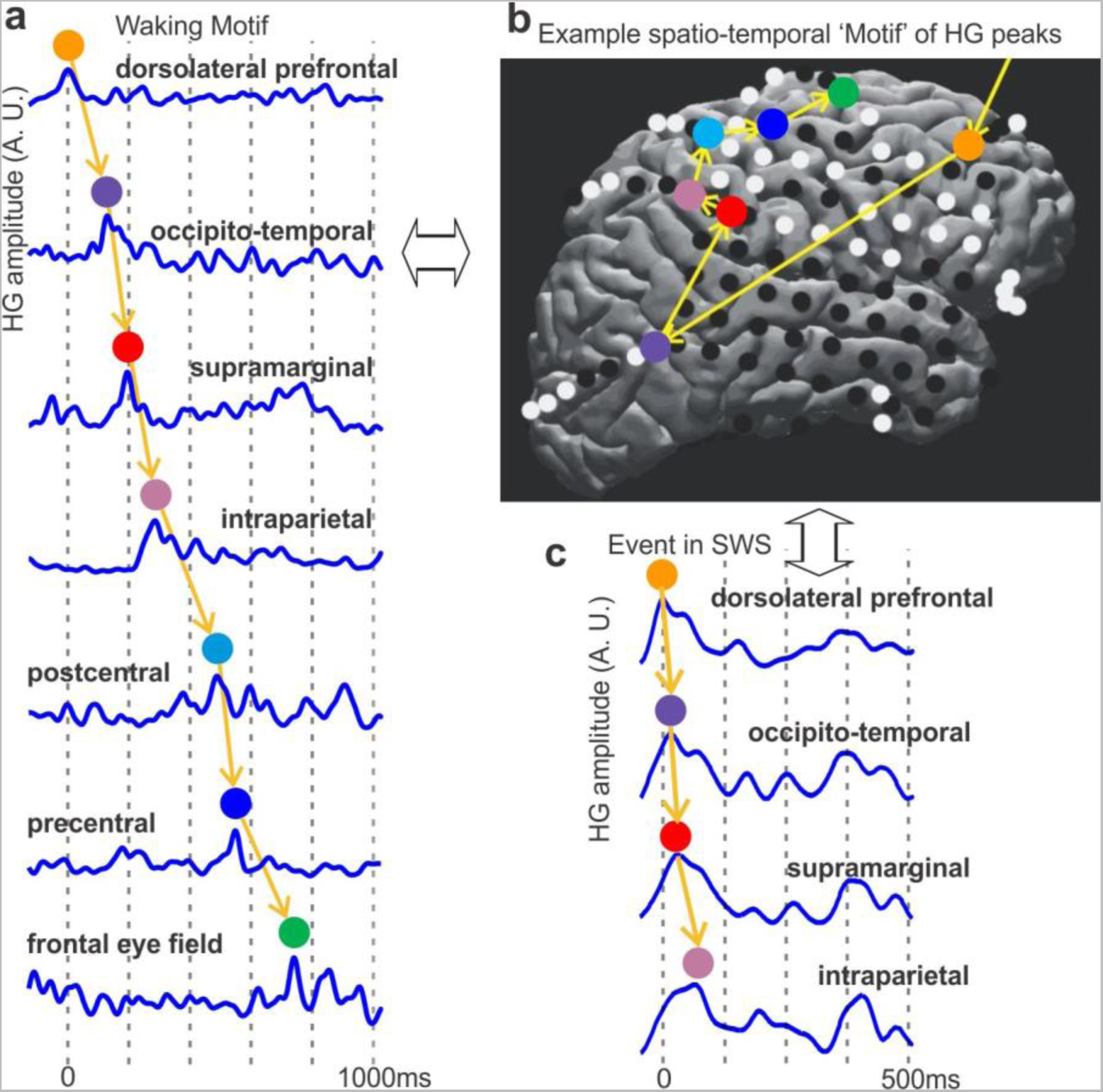
Spatio-temporal cortical activity Events and Motifs. **a**, Example Waking Event, comprising a sequence of HG peaks occurring within 2s. A.U.: arbitrary units. **b**, Example spatio-temporal Motif of HG peaks. Events were spatially- and temporally-clustered to obtain 60-250 representative Motifs for each subject, each with >20 Events (see “Motif selection and matching index analysis” in Methods and Fig. S1). Motifs with the most representative temporal order were matched against Events found in SWS (Fig. 3a). **c**, Example SWS Event whose HG peaks come from channels involved in the Motif to the left and are in the same temporal order as the same channels’ HG peaks in the Motif.

### Identification of consistent spatiotemporal Motifs of cortical activity during waking

For each subject, temporal sequences comprising HG peaks across multiple cortical locations (“Events”) (Supplementary Fig. S1, Fig. 1) were detected in both the single central waking period (W_0_) and the four SWS periods (Supplementary Fig. S2), two preceding W_0_ (thus comprising “Sleep-Pre”) and two following W_0_ (thus comprising “Sleep-Post”). Consistent Motifs of cortical activity were derived from waking Events using measures of similarity in the spatial distribution of active locations across the cortex, and the sequential order in which different cortical locations become active. Please note that we use the term “Event” to refer to a particular observed spatiotemporal sequence of HG peaks, and the term “Motif” to refer to the inferred consistent spatiotemporal pattern which could appear as several different Events with similar HG peak sequences. Our analysis detected Events and inferred Motifs during waking, then assimilated Events to Motifs during SWS in order to statistically compare their occurrence in different sleep periods. The spatial similarity of these Events was quantified using channel ensemble-wise correlation coefficients adopted from rodent studies of cortical patterns and memory replay ^34^. The temporal similarity of the Events was computed using matching indices ^4^, which required at least four distinct elements in a sequence; only Events containing at least 4 HG peaks from unique cortical loci were therefore involved in the identification of Motifs. Using these metrics, hierarchical clustering reduced the initial set of Events to a set of Motifs which consistently repeated during waking.

Based on the observed duration of cortical replay events from rodent data ^4^, we defined our Event selection criteria as having a total duration (i.e. time between the occurrence of the first and the last HG peak) of <2000 milliseconds, with no temporal overlap between any two Events. In order to test whether the occurrence of such Events is different from what would be expected under the null hypothesis of no consistent relationship in the activity of different cortical locations, the same Event detection procedure was applied to 1000 bootstrap replicates with shuffled inter-peak intervals. Because the HG peaks in different channels after shuffling no longer were temporally related to each other, the number of Events increased (Fig. 2a). However, because these Events no long resembled each other as often, the number of Motifs declined (Fig. 2b), and the number of channels contributing peaks to Events which were detected after shuffling was correspondingly smaller (Fig. 2c). These effects were all significant at the p<.001 level (i.e., all 1000 shuffles produced numbers outside the range of the actually observed data). This result held in all six subjects and also for each of the four SWS periods. These observations suggest that the detected Events represent non-random consistent spatio-temporal sequences of large peaks of cortical population firing.

### Recurrence of Motifs in the form of Motif-Event matches during sleep

These highly distributed cortical activity Motifs identified during waking were then matched with Events during slow wave sleep (SWS) in two preceding (“Sleep-Pre”) and two subsequent (“Sleep-Post”) nights. We concentrated our analysis on SWS rather than lighter NREM or REM sleep because it is characterized by the spindles and slow oscillations which are correlated with replay in rodents ^22,23^ and consolidation in humans ^25,35^. Stages N2 and N3 were combined because both contain spindles and slow oscillations (with delta power increasing and spindle power decreasing as N2 progresses into N3) ^36-38^. Also, combining N2 and N3 increased comparability to rodents who do not display that distinction during SWS ^26,39^. SWS was identified by increased low delta (< 2 Hz)/spindle (9-17 Hz) and decreased high gamma (70-190 Hz) power (Supplementary Fig. S3). Sleep depth was quantitatively matched using these characteristics in the Sleep-Pre vs. Sleep-Post epochs entered into analysis (Supplementary Fig. S3). Further confirmation of subject sleep/wake states was provided by state separation via PCA ^40^ (Supplementary Fig. S3).

**Figure 2.**
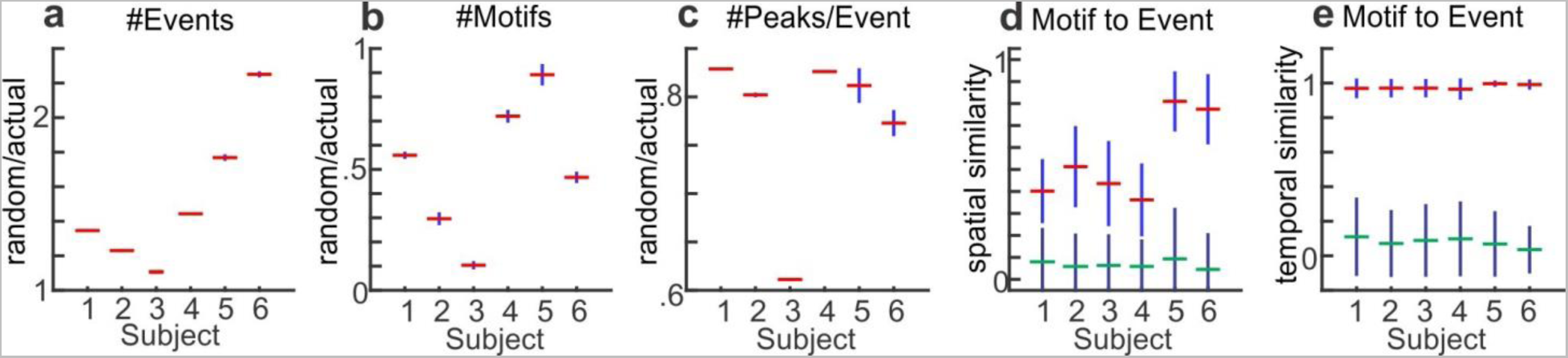
Waking Events and Motifs do not arise from chance, and are spatio-temporally similar. ***a***, The number of Events obtained from actual data is less than expected numbers from chance. Pseudo-Events were obtained from 1000 data shuffles (with the inter-peak intervals of each channel randomly permuted). For each subject, the distribution of ratios between the number of Events from shuffles and the actual number of Events is summarized. In this and all panels, the red horizontal line marks the mean, and the blue vertical line marks the standard deviation (SD). ***b***, The number of Motifs obtained from actual data is greater than expected numbers from chance. Pseudo-Motifs were obtained from >300 of the shuffles performed in ***a***. The distribution of ratios between the number of pseudo-Motifs from shuffles and the actual number of Motifs is summarized similar to ***a. c***. The number of channels involved in waking Events are thus greater than expected from chance. The distribution of ratios between the mean numbers of channels from pseudo-Events in ***a*** and the actual means are summarized as in ***a*** and ***b. d, e***: Events are more similar spatio-temporally to the Motifs they matched to than the other Motifs. ***d***, The distribution of spatial similarity indices calculated for Motif-to-Event matches (with red mean lines) and the distribution of indices calculated between Motifs and Events that are not matched to them (with green mean lines). The spatial index for each Motif-Event pair is defined as 2 × (number of channels in common)/(total number of channels in both the Motif and the Event). e, The distribution of temporal similarity indices for the Motif-Event pairs from ***d***. The temporal index for each Motif-Event pair is defined as [1+ (m-n)/(m+n)]/2, where m is the number of Event channel pairs in the same temporal order as the same channels in the Motif, and n is the number of such channel pairs in the opposite temporal order. For both ***d*** and ***e***, the blue distributions are significantly different from the black (two-sample Kolmogorov-Smirnov test, p < 0.0001), and the blue distributions have significantly greater medians (Mann-Whitney U test, p < 0.0001).

We identified Event matches to waking Motifs (Motif-Event matches) using the stringent criterion that all common channels must show gamma activity in the exact same temporal order, and sub-selected the Motifs under the consistency criteria, which remain unchanged for all subsequent analyses and are detailed as follows: we derived waking Motifs from the central waking period W_0_, and for each Motif, the number of Event matches were computed for S_-2_, S_-1_, S_1_, and S_2_ separately (Fig. 3a), with normalization to the total number of Events for each sleep period. However, if we denote the number of matches, or number of replaying frames (NRF) for a given sleep *n*, as NRF-S_*n*_, then for a given Motif to be considered for subsequent analyses, it needs to satisfy either of the following:

1. NRF-S_1_ > NRF-S_-1_, NRF-S_1_ > NRF-S_-2_, NRF-S_2_ > NRF-S_-1_, and NRF-S_2_ > NRF-S_-2_.
2. NRF-S_1_ < NRF-S_-1_, NRF-S_1_ < NRF-S_-2_, NRF-S_2_ < NRF-S_-1_, and NRF-S_2_ < NRF-S_-2_.

In particular, criterion 1 ensures that both NRFs from Sleep-Pre could not be greater than any NRFs from Sleep-Post for a given Motif to be considered “replaying”, and criterion 2 the opposite. In terms of both spatial and temporal similarity, the Events were more similar to their matched Motifs than to other Motifs with whom they did not match. In other words, for each subject, the distribution of spatial or temporal distance metrics for Motif-to-Event matches appeared to be significantly different (two-sample Kolmogorov-Smirnov test, p < 0.0001) and to have a significantly greater median (Mann-Whitney *U* test, p < 0.0001) when compared to the distribution of distances between unmatched Motifs and Events (Fig. 2d, e).

Our major finding was that in all six subjects (abbreviated as “Sub”), significantly more waking Motifs showed greater numbers of matches to activity in Sleep-Post than that in Sleep-Pre (Fig. 3c, p=0.0368 for Sub. 1, p=0.0022 for Sub. 2, p=0.0061 for Sub. 4, p<0.0001 for Sub. 3 and Sub. 6, p=0.0007 for Sub. 5). Thus, the majority of distributed spatiotemporal Motifs of cortical activity observed during waking occur more often in the following nights of SWS, as compared to the preceding nights. While replay sequences in rodents appear to be temporally compressed compared to waking events ^1,4^ the inter-peak intervals of our waking Motifs that recurred (i.e. had more Sleep-Post matches) were not significantly greater than those of the matching Events during SWS (Wilcoxon rank sum test, α=0.05) (Supplementary Fig. S4).

### Preferential re-activation during Sleep-Post is robust

We tested whether this finding of more Motifs with preferential re-activation during Sleep-Post was robust by varying our analysis in three ways. First, since replay Motifs during sleep (i.e. Events in sleep that matched waking Motifs) are not necessarily exactly identical to those during waking, we relaxed our criterion for a Motif-Event match. Rather than requiring that the activation order of all channels match exactly, we required only that the number of channel-pairs in the correct order was significantly above chance ^4^. Again, all subjects displayed a significant propensity to have Motifs with more Sleep-Post matches (Fig. 3d, p=0.3806 for Sub. 4, p=0.0048 for Sub. 1, p=0.0035 for Sub. 2, p=0.0036 for Sub. 5, p<0.0001 for Sub. 3 and Sub. 6). Second, we modified our criterion for considering a Motif to have occurred more often in Sleep-Post than Sleep-Pre periods. Rather than simply comparing the normalized number of Motif-Event matches in the two Sleep-Post periods to the number in the two Sleep-Pre periods, we required that the total number of matches exceed a threshold established by comparing Sleep-Pre and -Post Events to the same Motifs, whose channel orders were randomly shuffled ^4^. Only the four ECoG subjects had a sufficient number of Motifs to repeat our analysis with the stringent (Fig. 3e, p<0.0001 for each subject) and combinatorial requirements (Fig. 3f, p<0.0001 for each subject) after the shuffle tests. In all four subjects, more Motifs had a greater number of significant Sleep-Post matches than Sleep-Pre matches.

Third, we altered the criteria used to identify Events: we repeated our initial exact match analyses with different maximum Event duration in seconds (from 500 ms to 4000 ms, in 500 ms steps) and with different HG peak percentile thresholds (95^th^ to 99^th^ percentile, in 1 percentile steps). For all six subjects, we observed significant replay effect outside the initial parameter set of 99^th^ percentile, 2000 ms Event max duration (Supplementary Fig. S5). All subjects showed significant replay effect at the following parameter sets ([HG percentile, Event size]): [97^th^, 1 s], [98^th^, 1 s], [99^th^, 2 s], [99^th^, 2.5s]. Further, out of the 36 replay analyses with 99^th^ percentile threshold and max Event sizes from 1 s to 3.5 s (6 parameter sets, 6 subjects), 29 showed significant replay effect, and 0 showed significance in the opposite direction (i.e. majority of waking Motifs matching Sleep-Pre Events more often than Sleep-Post Events). Similarly, out of the 24 analyses with max Event size 1 s and percentile thresholds from 96^th^ to 99^th^, 20 showed significant replay effect, and 0 showed significance in the other direction. In summary, our finding that more spatiotemporal Motifs of HG activity observed during waking occur more often in Sleep-Post than in Sleep-Pre is robust to different methods for identifying matching HG Motifs, for judging differences in occurrence rate, and for identifying initial HG Events.

**Figure 3.**
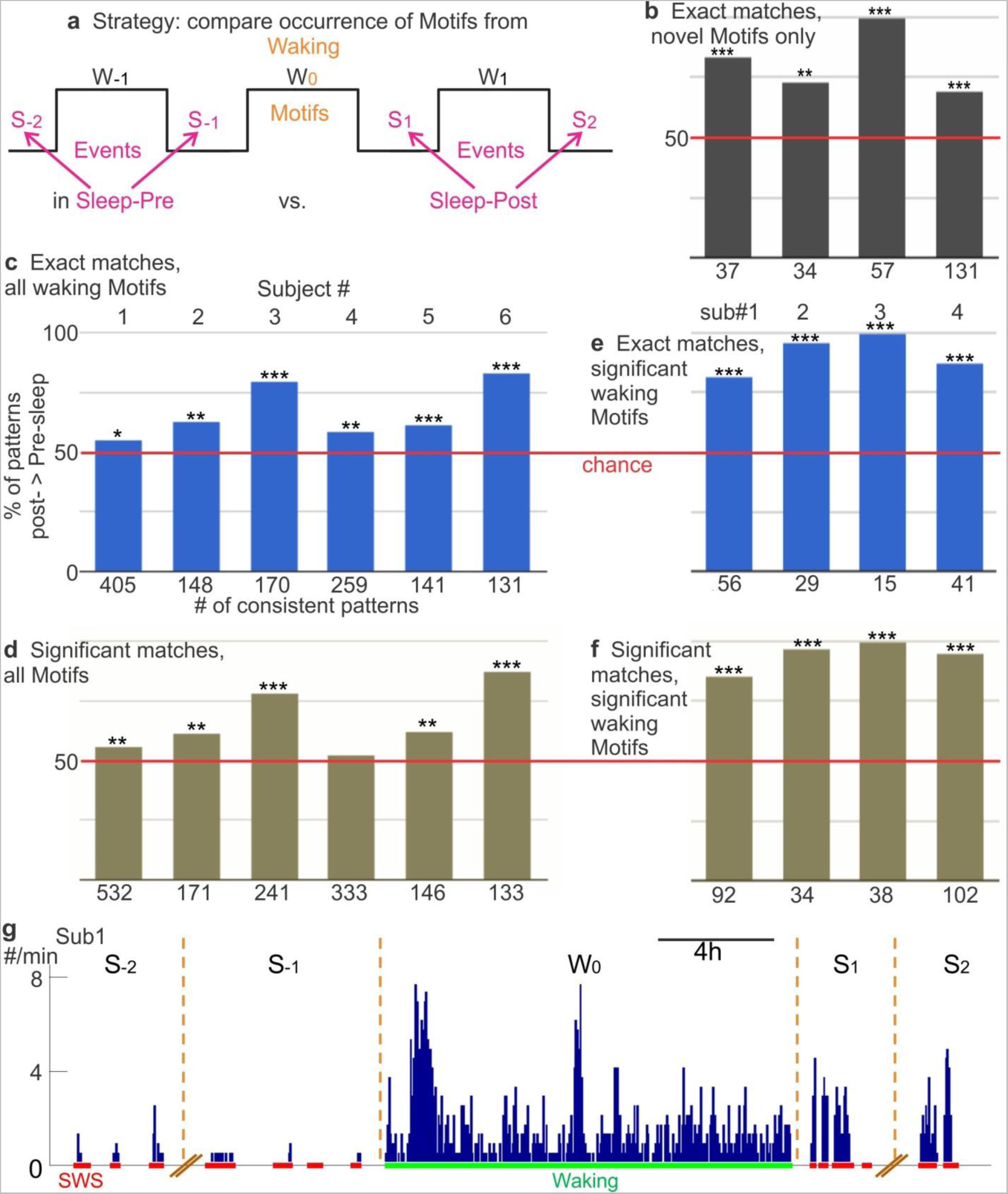
Events similar to waking cortical spatiotemporal activity Motifs occur more often in Sleep-Post than Sleep-Pre. ***a***, Schematic for Motif-Event matching. HG activity Motifs in the waking period W_0_ were matched to the Events found in Sleep-Pre (S_-2_, S_-1_) and Sleep-Post (S_1_, S_2_) periods. ***b-f***, Motif-Event matching results. Among waking Motifs with consistent (i.e. more Sleep-Pre than Sleep-Post matches for both S_-2_ and S_-1_, or *vice versa*) matches found across four nights, a higher percentage has more Sleep-Post matches than Sleep-Pre matches. The data are counts, and the null hypothesis is that the counts of Motifs showing greater occurrence in Sleep-Post than Sleep-Pre would be equal to those showing the converse, so a simple binomial test was performed on each subject. In ***b***, only Motifs found in W_0_ that did not occur in W_-1_ were used; for ***b, c*** and ***d***, exact matches were required; for ***e*** and ***f***, significant non-exact matches were also accepted; ***b, d*** and ***f*** only include waking Motifs that passed a significance test (see main text). SEEG subjects 5 and 6 did not have sufficient channels for the analyses in ***b, d*** and ***f***. Red lines represent chance (equal percentage of patterns with more Sleep-Post or more Sleep-Pre matches). (2-tailed binomial tests: ***b***, ^**^: p=0.009, ^***^:p<0.0001; ***c***, ^*^:p=0.0368, ^**^:p=0.0022 for 2, p=0.0061 for 4, ^***^:p<0.0001 for 3 and 6, p=0.0007 for 5; ***d***, ^***^:p<0.0001; ***e***, p=0.3806 for 4, ^**^:p=0.0048 for 1, p=0.0035 for 2, p=0.0036 for 5, ^***^: p<0.0001; ***f***, ^***^:p<0.0001). ***g***, The occurrence of exact matches across four nights (S) and one waking period (W) to waking Motifs that matched to more Events in Sleep-Post than Sleep-Pre. Histogram bin size: 8min.

As an additional control, we reversed sleep and waking periods and repeated our analyses. Specifically, we used identical parameters as in our initial analyses (exact match, 99^th^ percentile, 2000 ms max Event duration), but with a central sleep period as our source of Motifs, two preceding waking periods as the equivalent of “Sleep-Pre”, and two subsequent waking periods as the equivalent of “Sleep-Post”. In contrast to our original results, where all six subjects showed the replay effect, our control analyses yielded 3 subjects with no significant “replay” or “pre-play” (i.e. the majority of sleep “Motifs” matched more often to Events in preceding waking periods), two subjects with significant “replay” (p = 0.043 for Sub. 2, p < 0.0001 for Sub. 4), and one subject with significant “pre-play” (p = 0.01 for Sub. 1).

Since HG activity in epileptic patients is prone to contamination by interictal events, we eliminated >50% of the channels because they were located in the epileptogenic or adjacent zones (Table 1). For the remaining channels, we constructed an artifact detector based on wavelet decomposition (Supplementary Fig. S6) that was able to identify >99% of human expert hand-marked interictal events. Furthermore, the number of detected artifacts did not change significantly from Sleep-Pre to Sleep-Post (one-way ANOVA with each sleep as a “treatment group” and the percentage of interictal-containing 2000 ms time bins as the dependent variable, F-statistic: 2.493 on 3 and 1287 DF, p> 0.05), and indeed showed a slight nonsignificant decrease (median interictal rate across channels for Sleep-Pre was 5.41%, whereas the median rate for Sleep-Post was 4.96%), indicating that these events alone could not explain the increase in repeating Motifs (i.e. Motif-Event matches) from Sleep-Pre to Sleep-Post.

**Table 1.** List of subjects, age, sex, the types of electrodes used, the lengths of waking/sleep period recordings used, the median number of channels in an Event / Motif, the numbers of used (i.e. not dominated by epileptiform activity) versus available (i.e. the recorded voltage traces appear biological) cortical contacts, the number of waking Events / Motifs, and the overall occurrence rates of Motifs (i.e. Motif-to-Event matches). For subjects 5 and 6, only bipolar channels with clearly defined polarity inversions were kept for subsequent analyses. Hippocampal leads were too contaminated by epileptiform activity in subjects 1 and 4 to be used in this study. NA: not available. For a column with “/” separating numeric values and the column header, the numbers preceding the “/” symbol are described by the first part of this column’s header (i.e. the part that also precedes the “/” symbol), and the numbers following “/” are described by the second part of the header.

| Sub. ID | Age | Sex | Electrode Types | Waking Period Length (hours) | SWS Period Average Length (hours) | # of Channels Used / Available | Median # of Channels in Waking Event / Motif | Median # of Channels in Novel Waking Event / Motif | # of Waking Motifs / Novel Waking Motifs | # of Waking Events / Novel Waking Events | # of Waking Events Matched to Motif | # of Novel Waking Events Matched to Novel Motif | Median # of Channels in Waking / SWS Event Matched to Any Motif | Median # of Channels in Unmatched Waking Event | Median # of Channels in Unmatched SWS Event | Overall Rate for Waking Motifs (/h) | Overall Rate for Motifs in SWS (/h) |
| --- | --- | --- | --- | --- | --- | --- | --- | --- | --- | --- | --- | --- | --- | --- | --- | --- | --- |
| 11 | 27 | M | ECoG (grid+strip), SEEG | 19.6 | 2.1 ± 0.68 | 52 / 124 | 6 / 8 | 9 / 13 | 1677 / 116 | 14620 / 1731 | 4184 | 255 | 8 / 8 | 6 | 5 | 214 | 595 |
| 2 | 52 | F | ECoG (grid+strip), SEEG (hippocampus only) | 6.6 | 5.3 ± 1.9 | 44 / 96 | 6 / 7 | 5 / 9 | 411 / 67 | 3440 / 1277 | 1062 | 234 | 7 / 8 | 5 | 5 | 161 | 253 |
| 3 | 47 | M | ECoG (grid+strip) SEEG (hippocampus only) | 15.2 | 4.2 ± 0.83 | 65 / 124 | 7 / 9 | 5 / 10 | 481 / 146 | 4040 / 2080 | 1158 | 655 | 9 / 9 | 6 | 5 | 78 | 277 |
| 4 | 54 | F | ECoG (grid+strip), SEEG | 10.1 | 4.4 ± 1.2 | 75 / 121 | 7 / 9 | 7 / 15 | 774 / 350 | 8427 / 4285 | 1805 | 2289 | 10 / 11 | 7 | 5 | 184 | 590 |
| 5 | 53 | M | SEEG | 16.1 | 5.1 ± 0.62 | 12 / 80 | 4 / 4 | NA | 451 / NA | 2119 / NA | 1073 | NA | 4 / 5 | 4 | 4 | 67 | 58 |
| 6 | 24 | F | ECoG (strip), SEEG | 11.6 | 2.3 ± 1.0 | 18 / 116 | 4 / 5 | NA | 315 / NA | 2173 / NA | 689 | NA | 5 / 6 | 4 | 4 | 54 | 63 |
| 7 | 33 | F | ECoG (grid+strip) | 9.6 | 5.3 ± 1.5 | 60 / 126 | 6 / 8 | NA | 628 / NA | 6709 / NA | 1453 | NA | 8 / 9 | 5 | 5 | 98 | 433 |

### Novel Motifs are more likely to show greater Sleep-Post replay

Reasoning that novel experiences are more likely to require replay for consolidation, we attempted to restrict the analysis to novel Motifs by including only those that had not occurred in the previous waking period. With this requirement, the four ECoG subjects had significantly more novel Motifs (which passed the random shuffle test) with greater numbers of Sleep-Post than Sleep-Pre Motif-Event matches (Fig. 3b, with shuffle tests and exact matches, p=0.009 for Sub. 2, p < 0.0001 for others; SEEG subjects could not be examined because of too few qualifying Events due to low channel count). This result proves robust to different methods for identifying matching HG Motifs (i.e. allowing combinatorial matches and/or performing shuffle tests to evaluate Motifs) as well. Interestingly, more than 90% of the waking Motifs used in our initial analyses were actually novel in all four subjects.

**Figure 4.**
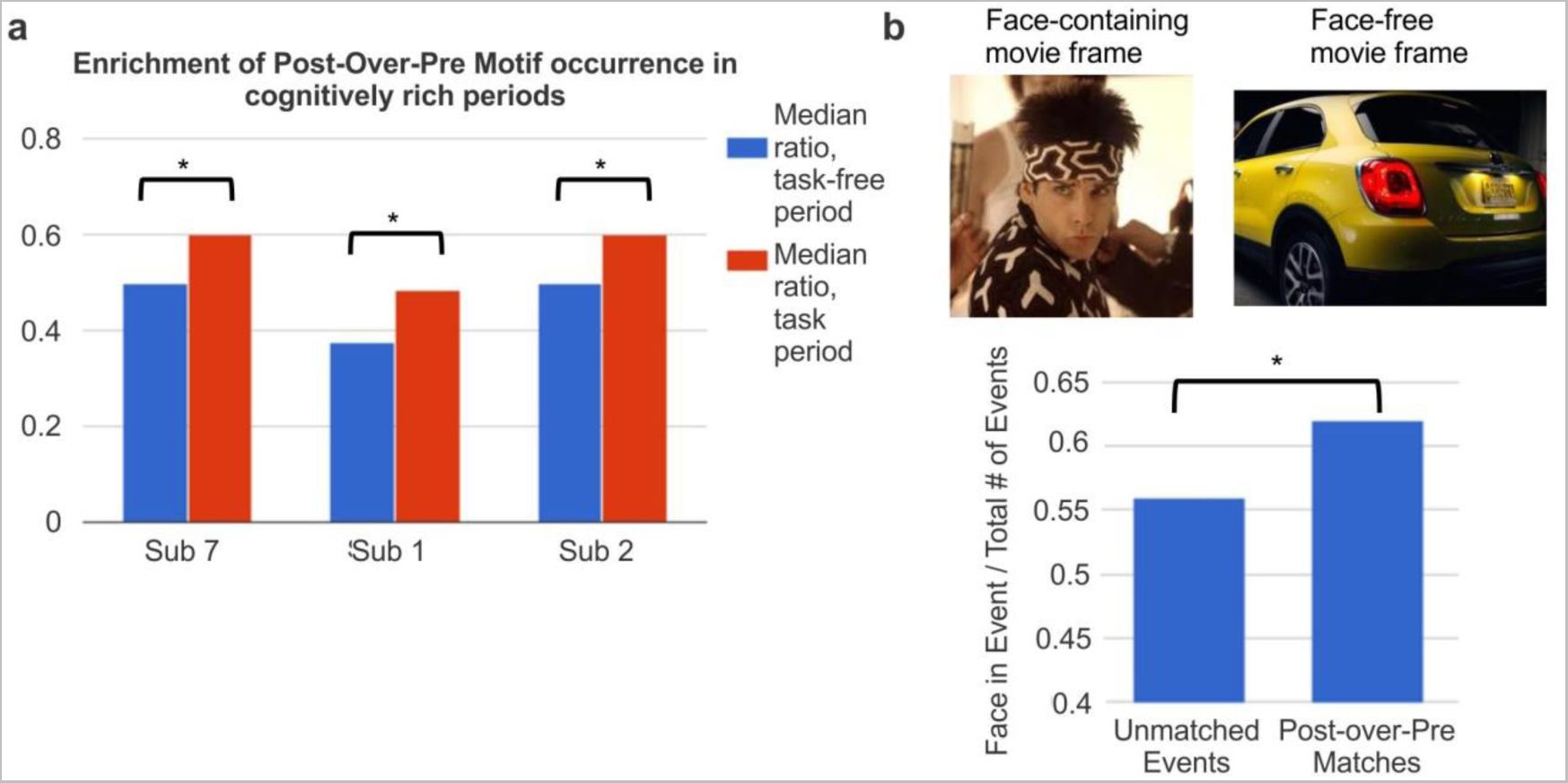
Motif occurrence enrichment in cognitively rich periods. ***a***, Motifs that matched more often to Events in Sleep-Post than to Events in Sleep-Pre (Post-over-Pre Motifs) preferentially occurred during task times in waking. For each randomly chosen 5-minute segment in waking, we calculated the following ratio: the number of Events similar to Post-over-Pre Motifs over the number of Events matched to all Motifs. 2000 random waking segments were selected, half within and half outside of the time for task performance. ^*^: p < 0.001 (Wilcoxon rank-sum test, alpha = 0.05). ***b***, Motifs whose occurrences (i.e. waking Event matches) overlapped with face-including movie frames were more likely than non-overlapping Motifs to have more matches in Sleep-Post. ^*^: two-tailed p-value (Chi-square test) = 0.0292.

### Enrichment of putative replay Motifs during cognitive activity

Waking period activity was recorded continuously during the uninterrupted clinical and personal activities typical of a patient in a hospital room under no acute distress. However, we were able to identify periods in 3 subjects when their mental engagement might be expected to be relatively greater due to cognitive testing or watching a movie. Thus, we tested if the HG Motifs with wake-sleep matches were related to cognitive activity in three subjects with information-rich waking time periods (>30 min) of cognitive task performance (including two of the subjects described above, and one additional subject not included in previous replay analyses because only data from one preceding and one subsequent sleep was available; see Supplementary Table S1). While the exact tasks performed differ across subjects because they were not specifically performed for the current study, we were able to identify task types and time periods based on experimental notes (see “Data collection” in Methods). Using a bootstrapping procedure (see “Statistical analysis for Events and Motifs” in Methods), we found that for all three subjects, Motifs occurring during information-rich periods were more likely than Motifs found elsewhere in waking to have more Motif-Event matches in Sleep-Post than in Sleep-Pre (p < 0.0001 for each subject, Wilcoxon rank sum test) (Fig. 4a). In addition to this apparent “enrichment” of putative replay by cognitive task performance, we also observed that for subject 7, who watched an entire movie, HG Motifs overlapping with face-including movie frames were more likely than non-overlapping Motifs to have more Motif-Event matches in Sleep-Post (2×2 chi-square test, α=0.05, p = 0.03) (Fig. 4b).

**Figure 5.**
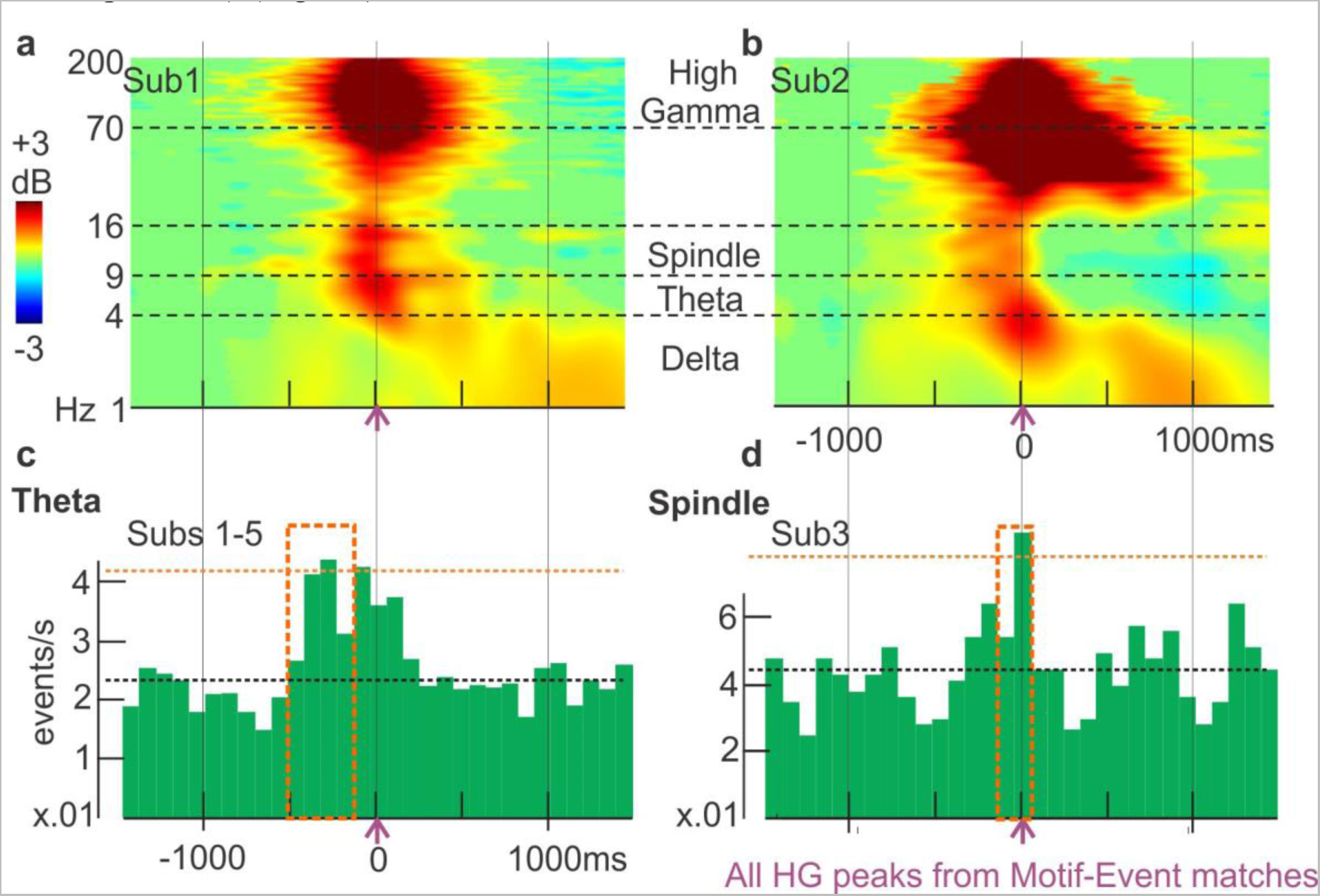
Putative replay frames are associated with spindle, delta, and theta activity. ***a-b***, Neocortical activity during sleep triggered on peaks of putative replay Events (i.e., Events matched to waking Motifs with “exact matches” criterion and no shuffle thresholds). Cortical theta, alpha, beta and gamma power increases for up to 1000 ms during putative replay events. Green mask for p>.05 (two-tailed bootstrap) from baseline period (−1500 ms to −1000 ms). Vertical solid lines / horizontal dash lines indicate shared X- or Y- axes across subplots, respectively. ***c-d***, Peri-HG-peak histograms of theta and spindle occurrences. Theta centers preferentially occur prior to Event peaks (Sub.1-5 grand average: p<0.001 from 600 ms before to 200 ms before Event HG peaks; FDR-corrected permutation tests), and spindles preferentially occur near Event peaks (Sub.2-3: p<0.001 in the 0.2s before the HG Event peaks; FDR-corrected permutation test). Orange boxes: significant time stretches. Putative replay frames are from the first Sleep-Post period. Similar effects were found for the second period (not shown). The black horizontal dash line indicates median ripple rate, and the orange horizontal dash line indicates the 99^th^ percentile.

### Relation of putative replay to cortical sleep spindles, theta bursts, and slow oscillations

In rats, replay by specific neuron ensembles occurs in the context of sharp wave-ripples in the hippocampus _29_;and spindles/delta activity in the neocortex ^22,23^. We thus examined delta power, theta oscillations, and spindle occurrences in cortical channels during the 3s surrounding putative replay Events in SWS, i.e. Events that matched Motifs with more Sleep-Post than Sleep-Pre matches (using “exact match” criterion, without applying shuffle thresholds). The HG amplitude peaks in these putative replay Events were generally associated with a ∼200 ms increase in broadband power surrounded by a prolonged increase in delta (Fig. 5a, b) in all subjects. In addition to elevated delta during putative replay Events, five of the six subjects displayed significant increases in theta and spindle occurrence in the period surrounding HG peaks from Motif-Event matches (the sixth was not included because too few spindles or theta bursts were detected, but showed similar trends). Theta occurrences consistently increased between 600 and 200 ms prior to the match peaks (p<0.001, using permutation tests with 1000 iterations, see “Time-frequency analysis and sleep graphoelement detection/analysis” in Methods). In contrast, the significant (p<0.001) increases in spindle likelihood could occur after 500 ms following the Motif-Event match peaks (in subjects 1, 4, 5), or in the 200 ms preceding the match peaks (in subjects 2, 3) (Fig. 5c, d).

**Figure 6.**
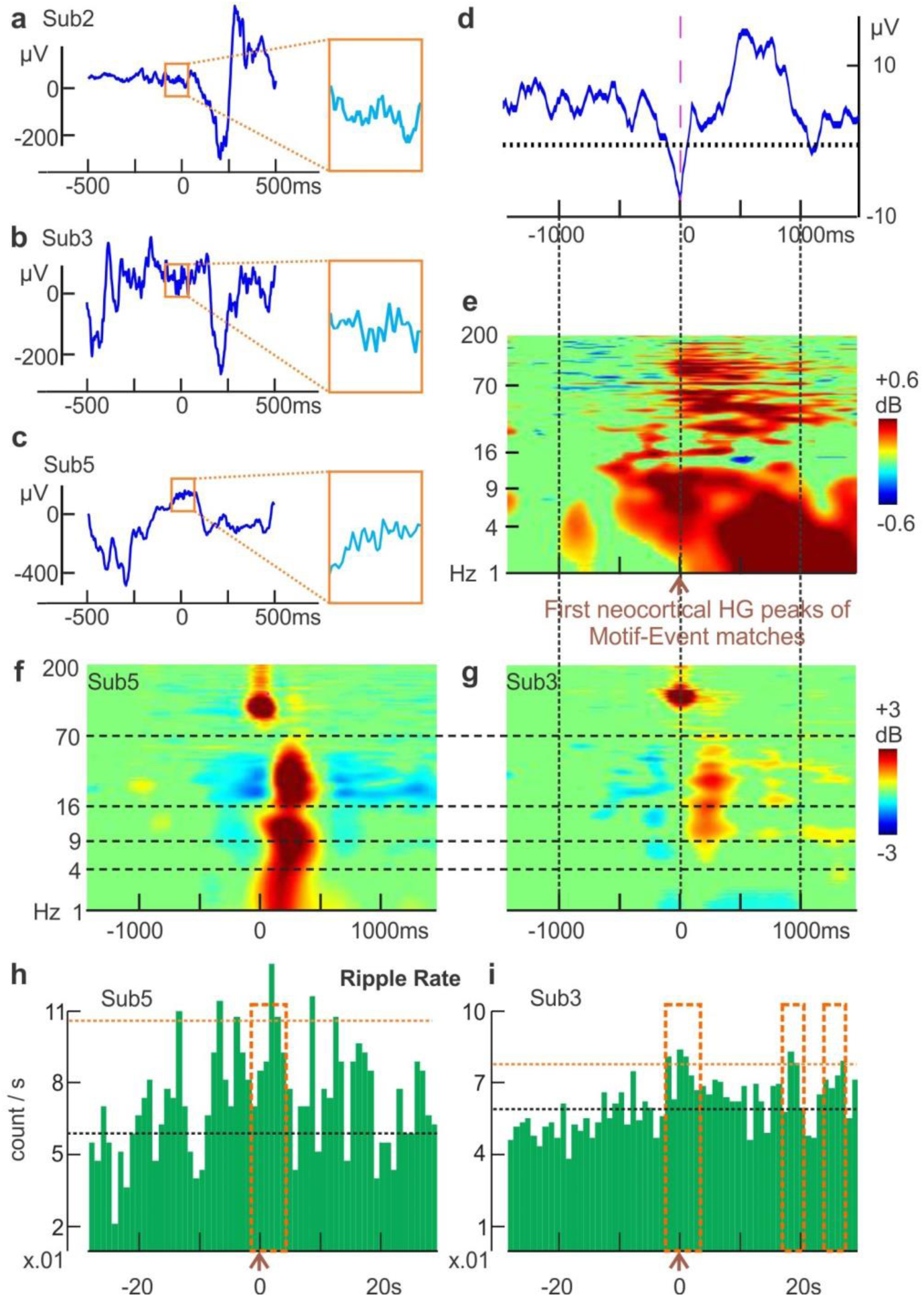
Putative replay frames are associated with hippocampal activity. ***a-c***, Examples of automatically detected hippocampal ripples (in orange squares, which covers 125 ms) and the surrounding LFP from subjects 2, 3, and 5, respectively. ***d***: Hippocampal LFP average triggered on 1^st^ cortical HG peaks of matched Events (exact matches, no shuffle thresholds). ***e***, Aggregate time-frequency plot with all three subjects (470 randomly selected trials each); broadband hippocampal activity increases for ∼1000 ms during matched Events. Power from 1-200 Hz (linear scale), with green mask for p > .05 (two-tailed bootstrap) from baseline period (−1500 ms to −1000 ms). ***f-g***, Time-frequency analysis of automatically detected ripples from posterior hippocampus shows power in the ripple frequency range (70-100 Hz). Vertical solid lines / horizontal dash lines that go across panels indicate shared X- or Y- axes across subplots, respectively. ***h-i***, Ripple occurrence rates become elevated near and after Motif-Event match peaks. The black horizontal dash line indicates median ripple rate, and the orange horizontal dash line indicates the 99^th^ percentile. Time windows that show significance are described here as [start,end] in milliseconds, where negative/positive values refer to the temporal distance from trigger times: p=0.039 at [-2000,2000] and p<0.001 at [0,4000] for Sub.5; p=0.013 at [-2000,2000], p=0.0446 at [0,4000], p=0.039 at [1600,2000] and at [2400,2800] for Sub.3 (FDR-corrected permutation tests). Orange boxes: significant time stretches. Due to the long time-base, only the first HG peak from a given Event would be used in this analysis. All Events are from the first Sleep-Post period. Similar effects were found for the second period (not shown).

### Relation of putative replay Events to hippocampal activity

Simultaneous cortical ECoG and hippocampal SEEG recordings were available in three of our six subjects (Table 1), where we detected hippocampal sharp-wave ripples (Fig. 6a-c). Using the first (among the channels that matched the waking templates) HG peaks of putative cortical replay Events as triggers (Events were obtained under “exact match” criterion, without applying shuffle thresholds), a sharp transient was noted in the average hippocampal LFP (Fig. 6d). In all three subjects, posterior hippocampal LFP showed slow activity coincident with significant broadband activation including frequencies over 100 Hz (Fig. 6e). This frequency range could reflect both ripples and unit firing. In rats, hippocampal sharp waves coincide with cortical transitions to up-states during SWS ^41^, and with coordinated replay in both cortex and hippocampus ^4^ In our study, the prolonged (∼1000 ms) increase in hippocampal HG suggests continuing hippocampal-cortical interaction as the successive cortical nodes in a Motif-Event match are activated. Indeed, in all three subjects (Sub. 2, 3, 5) with hippocampal contacts (left and right posterior hippocampi), morphologically typical sharp-wave ripples were recorded (but Sub. 2 had 10 times less than the others), with similar frequency profiles to previously reported primate ^42^ and human ^37^ ripples (Fig. 6f, g). In both Sub. 3 and Sub. 5, we observed a significant prolonged (up to at least 4s post-trigger) elevation in ripple occurrence rates after Motif-Event match peaks (Fig. 6h, i; p=0.039 at [-2s,2s] and p<0.001 at [0,4s] for Sub. 5; p=0.013 at [-2s,2s], p=0.0446 at [0,4s], p=0.039 at [16s,20s] and at [24s,28s] for Sub. 3).

### Selective involvement of cortical areas in putative replay Events

Visual inspection suggests that some cortical locations were involved more frequently in putative replay Events, and/or selectively at the onset of such Events (Fig. 7). We tested in subjects 1-6 whether the amount of HG peaks (which are from both waking and SWS Events that matched to waking Motifs with higher occurrence in Sleep-Post than in Sleep-Pre) found at each cortical location was significantly different from chance by a modified binomial test (p < 0.05 with FDR correction, see “Statistical analysis for Events and Motifs” in Methods). For all subjects, more than 50% of channels showed a significant difference, either greater or less than expected under the null hypothesis that all channels contributed equally to motifs. Similarly, the likelihood for each cortical location to have high gamma peaks in the beginning of an Event was found to be significantly different than chance for 20-80% of each subject’s channels (permutation tests, p < 0.05 with FDR correction). Despite this evidence of nonrandom distribution of sites with more or less involvement in Motifs, earlier or later in the sequence, there was no obvious anatomical focus consistent across all subjects. However, in subjects 1, 2, and possibly 4, there seemed to be a tendency for the parietal-temporo-occipital junctions to have more HG peaks than expected by chance.

**Figure 7.**
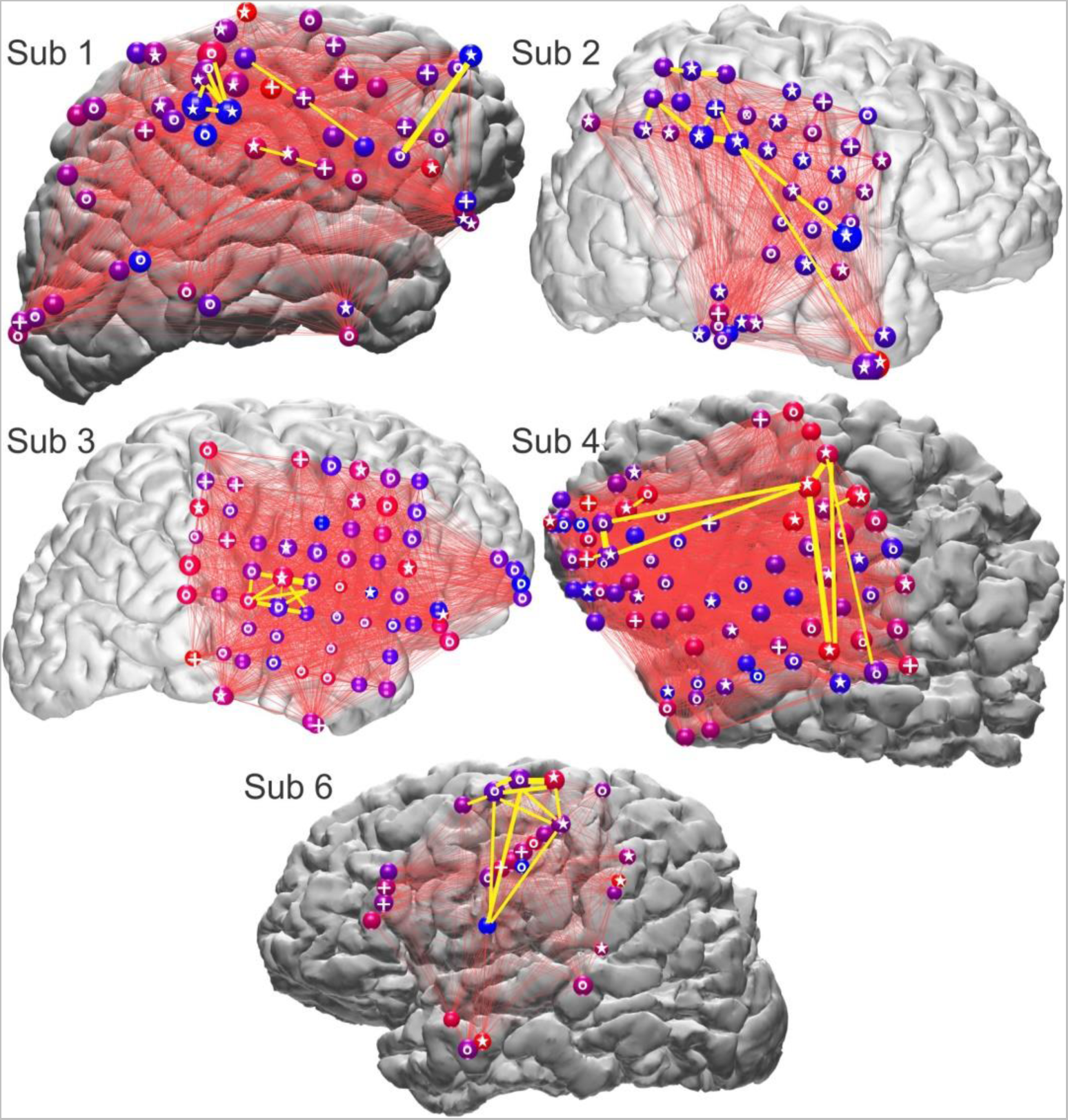
Participation of widespread cortical areas in Motifs. Spheres represent ECoG electrodes, with circles (o), plusses (+), or stars (☆) on the spheres indicating that they showed a significant deviation from chance (FDR corrected p<.05) in their occurrence rate, in their temporal order in the motifs, or in both, respectively. Specifically, a circle indicates that the amount of high gamma peaks seen at this cortical location was significantly different (more or less) from expected value based on chance alone. Similarly, a plus indicates that the likelihood for this cortical location to have high gamma peaks in the first quarter of an Event was significantly different (higher or lower) from chance, i.e. 25%. The size of each sphere indicates HG peak occurrence rate; color indicates the likelihood of HG peaks being early versus late in an Event (red: early, blue: late). Thin red lines connect the electrode pairs that produced temporally adjacent HG peaks within Events. Yellow lines connect the ten electrode pairs that produced the largest numbers of temporally adjacent HG peaks. Plotted are HG peaks from Motifs occurring in all waking and sleep periods (i.e. from both waking and SWS (the two Sleep-Post periods) Events that matched to waking Motifs with higher occurrence in Sleep-Post than in Sleep-Pre).

## Discussion

We present the first clear demonstration of locally-recorded cortical replay in humans, in the form of large-scale spatio-temporal HG activity identified during waking and recurring during subsequent sleep periods. Each Motif typically involved widespread sensory, motor and association areas. While consistent with some primate recordings ^8^ and human hemodynamic studies ^30^, the current study is the first to our knowledge showing that both the initial waking representation and the subsequent sleep replay could involve widespread dynamic sequences using intracranial electrophysiology.

In order to demonstrate replay of widespread cortical spatio-temporal activity patterns, it was necessary to first demonstrate them in our recordings. We used clustering and statistical methods adapted from unit recordings studies in rodents to identify consistent spatiotemporal patterns of HG peaks (‘Motifs’), and then tested if they were different from chance by comparing them to Motifs found when the temporal relationship between HG peaks was disrupted with randomization. Randomization resulted in a highly significant decrease in the number and size of Motifs were found. While consistent spatio-temporal patterns have been observed to occur spontaneously with hemodynamic measures over widespread cortical areas ^43^, and locally using multi-microelectrode recordings ^34,44^, we are not aware of their having previously demonstrated in human cortical electrophysiological recordings. When the intervals between HG peaks were randomized fewer and smaller Motifs were found. This finding of highly distributed recurring spatiotemporal activity patterns confirms a basic assumption of cortical processing models ^45^.

Our finding of Motif replay was robust to different statistical criteria for identifying Events in sleep that matched waking Motifs, for determining if Motif occurrence rate differed between Sleep-Pre and Sleep-Post periods, and when only novel Motifs were considered. In fact, most of the recurring Motifs had not occurred in previous waking periods. Furthermore, preferential replay in Sleep-Post were generally replicated when the maximum pattern durations ranging from 1000ms to 3000 ms (rather than the 2000 ms we initially used) and HG peak selection from 97 to 99th percentiles (Supplementary Fig. S5). As an additional control, no consistent results were obtained when we reversed sleep and waking periods and repeated our analyses. The electrophysiological context of recurring Motifs is thus consistent with those reported in rodents and humans to be associated with replay and/or consolidation. Notably, recurring Motifs (i.e. sleep Events that matched to waking Motifs) were associated with increased cortical sleep spindle and delta activity which are associated with not only unit firing replay in rodents ^22,23^ but also memory consolidation in humans ^25,28,35^. Critically, the putative replay Events in our study were accompanied by strong sustained hippocampal activation, resembling sharp waves and ripples. Hippocampal sharp wave-ripples have a strong relationship to hippocampal unit replay in rodents ^4^, and presumably that is also true in humans. Thus, their association with cortical Motifs provides a neurophysiological mechanism for the hypothesized temporary hippocampal trace to contribute to the construction of the recurring cortical Motif.

We observed that putative replay continued for at least 2 nights after the waking period in which the Motif was originally detected (Fig. 3, Supplementary Fig. S7). This is consistent with the prolonged retrograde amnesia that occurs after hippocampal damage, suggesting an extended consolidation period ^46^. The putative replay events lasted about 1000-2000 milliseconds in humans, consistent with rodent data ^4^. This duration may reflect the algorithm used to identify spatiotemporal Motifs; longer or shorter Motifs could be found when we altered our maximum Event duration (Supplementary Fig. S5). Unlike rodents, the replaying Motifs we observed did not appear to be accelerated compared to their waking occurrence. This could reflect differences in approach and/or actual species differences. Humans typically live about 70 years and have one long sleep period every night. Rats typically live about 2 years and have many sleep periods throughout the day, staying in each state for as little as a few minutes ^47^. Furthermore, compared to rats, humans have about 30 times more neocortical cells per hippocampal pyramidal neuron ^48–51^. Thus, hippocampal orchestration of neocortical replay may be modified in humans, and the characteristics of replay occurrence may therefore differ substantially.

Strong circumstantial evidence supports a key role for replay during sleep for memory consolidation in rodents. For example, the reactivation of spatial goal-related firing during SWRs after learning predicts later memory recall, an effect which is depends on NMDA-mediated reactivation of goal-related firing pattern reactivations during ripples ^3^. In addition, biasing behavior with external modulation of replay has been demonstrated by pairing spontaneous firing of hippocampal place cells with reward signal ^7^, as well as by auditory cueing of spatial preference during NREM sleep ^52^. In our study, recurring Events during SWS showed a strong preference to arise from more cognitively rich periods in waking, but further work is needed to determine their experiential content, which would provide the leverage to obtain direct behavioral evidence that their replay is associated with memory consolidation.

We demonstrate here replay during sleep in humans, and its similarity in certain key features to replay in rodents (occurrence in SWS, close relation to hippocampal activity and to cortical sleep graphoelements), but its role in memory consolidation remains unexplored. Regardless of their relationship to memory consolidation, our results demonstrate the existence of consistent widespread spatiotemporal patterns of cortical population firing. Such patterns are fundamental to associationist perspectives on cortical function ^53^, but have not previously been described in humans.

## Materials and Methods

### Data collection

Seven subjects (3 male, 4 female) with long-standing drug-resistant partial seizures underwent ECoG macro-electrode (subdural grid and strip) and/or SEEG depth electrode implantation (4 with ECoG and limited SEEG, 2 with SEEG only, 1 with ECoG only). Subjects were all right-handed except for patient 5, who was ambidextrous. For patients 2-5 and 7, drugs were either no longer given to the patients by the time the recordings used in analysis had begun, or were provided in exactly the same dosage every day analyzed. Subjects 1 and 6 had changes to their medical regimen after the day of the first Sleep-Pre, but not for the other three days used in this study: subject 1 discontinued 200 mg oral carbamazepine and subject 6 started 50 mg oral quetiapine. For subjects 1, 3, 5, 6, and 7, quantitative clinical neuropsychology information was available, and subjects 2 and 4 were noted by clinicians to be cognitively normal. Subjects 1 and 7 had average scores on the Wechsler Adult Intelligence Scale (WAIS-IV) in Verbal Comprehension Index (Sub. 1: 100, Sub. 7: 105) and Perceptual Organization Index (Sub. 1: 107, Sub. 7: 100). Subject 6, who had lower scores (∼80) on WAIS-IV, was a native Arabic speaker who did not speak English fluently. Subjects 3 and 4 received evaluation on WAIS-III: subject 3 had average (86) Verbal IQ, high (124) Performance IQ, and average Wechsler Memory Scale (WMS) recent memory test scores; subject 4 had low average scores (74/79) in Verbal/Performance IQ and in WMS.

Electrode targets and implantation duration were chosen entirely on clinical grounds. All methods were carried out in accordance with clinical guidelines and regulations at Comprehensive Epilepsy Center (New York University School of Medicine) and at Massachusetts General Hospital. All subjects gave fully informed consent for data usage. All experiment protocols were monitored and approved by the Institutional Review Boards at New York University and Partners HealthCare. After implantation, electrodes were located using CT and MRI ^54,55^. Continuous recordings from grid and strip macro-electrode arrays were made over the course of clinical monitoring for spontaneous seizures, with sampling rates at 500 or 512 Hz, using 128-channel clinical cable telemetry systems. All SEEG contacts were converted to a bipolar montage. Each pair of contacts (separated by 3.5 mm center-to-center) was only included if one contact was near the pial surface and the other near the white-to-gray transition, according to preoperative MRI aligned with postoperative CT ^56^, confirmed with neurophysiological criteria (high amplitude, coherence and inversion of spontaneous activity between the contacts). These bipolar transcortical derivations ensure that activity is locally generated ^38^. During part of the recording periods, subjects 1, 2, and 7 engaged in cognitive activities (>30 min on one given day of their stays) as research participation for other groups; the patients’ spontaneous waking behaviors were not interfered with otherwise. Subject 1 watched movie trailers (full video/audio) and performed a name-face association task, subject 2 performed an auditory pattern detection task and an auditory sequence discrimination task, and subject 7 watched the movie “Zoolander” (full video/audio) without interruption. Detailed task setup/descriptions and movie processing steps can be found in the Supplementary Methods.

### Preprocessing and SWS selection

Recordings were anonymized and converted into the European Data Format (EDF). Subsequent data preprocessing was performed in MATLAB; the Fieldtrip toolbox (RRID:SCR_004849) ^57^ was used for bandpass filters, line noise removal, and visual inspection. High gamma amplitude (HG) over time was estimated by smoothing the analytic amplitude of 70-190 Hz bandpassed data with a 100 ms 1D Gaussian kernel (Supplementary Fig. S1). Raw data corresponding to all SWS periods in each sleep were concatenated prior to HG peak detection. The top 1 percent of all local HG maxima for each channel was used for subsequent analyses. Given the lack of access to clinical hypnograms, patient monitoring videos, scalp EEG, electrooculogram, or any form of sleep staging hand marks for most subjects, instead of following the popular AAMS manual criteria ^58^, SWS periods (as a combination of NREM sleep stages N2 and N3, where N2 and N3 forms a temporal continuum with progressively increased slow oscillation content) were identified by prolonged (> 5 min) elevations in low delta band (< 2 Hz) activity amplitude and corresponding declines in HG and initially high but gradually decreasing spindle band (9-17 Hz) amplitudes (Supplementary Fig. S3). Each SWS period was then verified by visual inspection for the presence of sleep graphoelements. Sleep-wake boundaries were defined as 1 hour before the first in a series of SWS periods (which are considered as in the same “sleep” if less than 2 hours apart), or 1 hour after the last in a series. Time segments between bouts of SWS and with no sleep graphoelements were considered REM or transient waking, and were removed from analyses.

Confirmation of subject sleep/wake states was provided by methods adapted from an early rodent behavioral state separation study ^40^, where periods of wakefulness and different sleep stages were identified as clusters on 2D histograms of the first principal components (PCs) for two frequency ratios, computed from Fourier transform integrals over different frequency bands.

For the current analysis, these methods were modified as follows: one PC was computed from the ratio of the Fourier transform integral for 0.5-16 Hz over the integral for 0.5-55 Hz, and the other PC from the ratio of the 0.5-3 Hz integral over the 0.5-16 Hz integral. This method was validated against manual classification of sleep stages in subjects 2 and 3, by experienced clinicians (Supplementary Fig. S3). To ensure that the depth of sleep is equal for Sleep-Pre and Sleep-Post, normalized differences were calculated from the channel-wise z-scored differences between Sleep-Pre and Sleep-Post delta amplitudes (sum of two Sleep-Pre SWS periods minus sum of two Sleep-Post SWS periods) for each subject, and evaluated via one-sample sign tests. Randomly centered 1 second time bins with greater than or less than median delta amplitude were discarded iteratively from Sleep-Pre and Sleep-Post as needed, until no significant difference between Sleep-Pre and Sleep-Post remained (Supplementary Fig. S3).

### Artifact rejection

Since HG peaks may sometimes arise from artifacts or epileptiform activity, prior to bandpassing signal over the HG band, 2000 milliseconds of data centered around each HG peak underwent 1-D wavelet (Haar and Daubechies) decomposition for the detection and removal of sharp transients (i.e. signal discontinuities) that might contaminate the HG band. In addition, HG peaks that occurred in close temporal proximity (< 100 ms) across more than 20% of all channels were discarded. Within each channel, HG peaks whose amplitudes exceeded 20 standard deviations above the mean were also excluded, and the minimum distance between same-channel HG peaks eligible for downstream analysis was set to 100 ms. Channels were not analyzed if the local cortex consistently displayed or was marked by clinicians as prone to interictal (or otherwise abnormal) activity (Table 1).

### Event selection

HG peak Events in the form of vectors with integers designating different channels were created as follows: 2000-millisecond sliding windows were moved across the 4 SWS periods (two preceding the waking period, two subsequent) and the waking period data in 50 ms increments, with each movement yielding a different number of outlier HG peaks within the current window (Supplementary Fig. S1). The windows that yielded local maxima of HG peak numbers were taken as Events. A minimum of 4 peaks was required for an activity Event.

In the analysis limited to novel Events, the same initial procedure was followed except that the requirement that the Event represent a local maximum of HG peak numbers was relaxed to increase the sensitivity to all patterns that might recur. Instead, Events were defined by using each HG peak outlier as the starting point, and extending the Event to include subsequent peaks unless the inclusion of another peak would increase the Event length beyond 2000 milliseconds. The novel Event search was therefore not performed with a sliding window, but with a “saltatory” window (with a maximum length of 2000 milliseconds) that starts at the first (in temporal order) HG peak in the data, reaches for up to 2000 milliseconds to include subsequent HG peaks, terminates, and restarts at the next HG peak in the data. This search procedure would be terminated either when all HG peaks belong to some elastic window, or when the last saltatory window would contain less than 4 HG peaks.

In all analyses, each Event was then binarized (0 for no HG in a channel, 1 for having at least one HG in a channel), and converted further into a 2-by-n spatio-temporal representation (one column for time indices, the other for channel number), where n is the number of total HG peaks in the pattern (Supplementary Fig. S1). In cases where a channel had more than one HG outlier peak within the same Event, the multiple peaks from this channel were replaced with a single peak located at the median position in time for all peaks from this channel, so that each Event contains only one HG peak per channel.

### Motif selection and matching index analysis

Spatiotemporal activity Motifs were selected via metrics derived from rat cortical electrophysiology studies as follows: hierarchical clustering of the waking period Events was performed in R (with the packages Fast Hierarchical Clustering Routine ^59^ and Dynamic Tree Cut ^60^ using Phi-coefficient-derived distance matrices ^34^ and Ward’s method ^59^ to group Events according to spatial distribution similarity. To select waking Motifs by further grouping Events according to HG peak order, and to match waking Motifs to SWS Events, temporal order similarity for waking Events in each cluster was then obtained via the matching index (MI) algorithm, which was implemented (using HG peaks in place of spike train convolutions) as previously described ^4^ (in MATLAB with custom scripts and in R with the TraMineR toolbox ^61^), with the additional heuristic criterion that for each comparison, only the channels in common were used to calculate MI. Note that our measure of pattern similarity requires that each spatial location has a unique position in the temporal sequence, since this measure relies on counting the number of unit pairs (ECoG channel pairs in our case) in the same versus opposite order from a given Motif-Event match ^4^. We therefore combined multiple HG peaks from the same channel in a given Event to a single peak at the median position in time of all HG peaks from that channel. This is a potential drawback in terms of Motif-Event match fidelity, as multiple HG peaks from the same channel in a stereotypical temporal order might be a key aspect of a given recurring Motif. However, if true, such stereotypy across multiple iterations/Events may also lead to similar median peak positions for that particular channel, relative to HG peak positions from other channels.

Based on the MI outcome (a distance matrix of match probabilities), each cluster of spatially similar waking Events was split into subclusters. Representative Motifs were chosen as those that include 90% of a given subcluster’s Events in their neighborhoods, which were defined as 20% of the maximum distance between two Events for a given subcluster. These were then matched with MI to the Events in SWS, and each representative was associated with a certain number of replay frames (i.e. sleep Events that matched the waking Motif, or “Motif-Event matches”) for each SWS period. If multiple Motifs were matched to the same SWS Event, for statistical purposes, the number of replay frames associated with each waking Motif for that night was incremented by 1/N instead of 1, where N equals the number of waking Motifs that matched to this common Event. So, if there were exactly three Motifs matched to the same Event, each Motif would have its number of replay frames incremented by only a third. Thus, the total count increments from any given Event would equal exactly 1 regardless of how many Motifs it matched, as long as it matched at least one. The resulting replay frame numbers for each SWS period were then normalized by the total number of Events found in that SWS period. For the random shuffle test of replay frame number significance ^4^, the two subjects with only SEEG contacts used for matching were not examined, since only Motif-Event matches with >6 channels satisfy the test’s requirement, and few Motif-Event matches from these subjects exceed 6 channels in length.

Note that our analysis did not search for repetition during sleep of Events which occurred only once during waking. We required that Motifs were matched during waking by at least 2 Events, and such Events only comprised ∼30% of waking Events on average across subjects. The possibility that these unique activity patterns also replay during subsequent sleep was not explored in the current study because it appeared prudent for us to perform our analyses on only the consistently recurring patterns that would be more likely to reflect reactivation of waking experiences. One requirement of such an investigation would be to determine if the unique Events are significantly non-random. In rodents, hippocampal replay was observed not only in SWS, but also during waking ^62^; in humans, fMRI evidence suggests that event-specific reactivation, which predicts later successful recall, occurs in both hippocampus and cortex spontaneously after learning during waking ^12^. Thus, while we could not exclude the possibility that rare activity patterns might also be involved in consolidation, we refrained from performing our novel analyses on unique, one-off patterns.

### Statistical analyses for Events and Motifs

All statistical analyses involving hypothesis testing described under this section and the following section (LFP analysis) have α = 0.05. Putative replay of waking Motifs was statistically evaluated as follows. First, Motifs were counted which had more Motif-Event matches during each of the Sleep-Pres (S_-2_ and S_-1_) than in either of the Sleep-Posts (S_1_ and S_2_). This was compared to the number with the opposite pattern (more matches during each of the Sleep-Posts than in either of the Sleep-Pres), using 2-tailed binomial tests (hypothesized success rate set to 0.5). In order to test the sensitivity of the analysis to the choices described above regarding inclusion of Motifs and criteria for Motif-Event matches, binomial tests were also performed after supplementary analyses with the following restrictions that produced different numbers of matches or different sets of Motifs. First, we required the common cortical loci in matching Events and Motifs to have the exact same HG peak temporal order (“stringent”) or not (“combinatorial”). Second, following Ji and Wilson ^4^, we used only Motifs whose total number of Motif-Event matches across all nights exceeded Motif-wise thresholds established by matching Sleep-Pre and -Post Events to the same Motifs with their channel order randomly shuffled 1000 times. Third, we used only novel Motifs.

In order to test if the numbers of waking/sleep Events observed, the mean number of HG peaks in Events, and the kurtosis of HG peak number distributions were greater than expected from the null hypothesis that HG peaks occur randomly across the cortex, we compared the number of observed Events (with maximum duration of 2000 ms and at least 4 unique cortical locations), mean HG peak numbers, and kurtosis values from the original data to the numbers derived from 1000 sets of shuffled peak locations, where the total number of HG peaks and the location of the first HG peak from each channel remained the same as in the original, but the inter-peak intervals were shuffled randomly in temporal order over the same waking/sleep periods. The threshold for bootstrapping significance was set at 0.05.

In order to test if the task periods were enriched in waking Motifs that had more Motif-Event matches in Sleep-Post than in Sleep-Pre (note that, unlike previous replay analyses, only one Sleep-Post and one Sleep-Pre periods were used), we performed the following bootstrapping procedure: template-matching analyses were performed with one Sleep-Pre and one Sleep-Post per subject, using 10,000 random (allowing overlap) 5-minute time bins selected from either task or no-task waking periods. We then computed for each time bin a ratio between the number of waking Motifs with more Sleep-Post matches and the total number of Motifs with at least one match in sleep. These ratios were compared between task-on and task-off periods using the Wilcoxon signed-rank test.

In order to test if a given cortical location produced significantly more or less HG peaks than expected from chance, we took all the waking Motifs with more Motif-Event matches in Sleep-Post and the sleep Motifs they matched to and obtained, for each channel, the numbers of Motifs involving (“successes”) and not involving (“failures”) that channel. Each success-failure ratio was then evaluated via binomial test, with the expected success probability set as follows: for each subject, the mean number of HG peaks involved in a Motif-Event match was divided by the total number of channels. Thus, the null hypothesis was that all channels contributed equally to all motifs. Similarly, in order to test if a given cortical location was significantly more or less likely to produce HG peaks early (i.e. the first 1/4 of all channels involved), the number of successes and failures obtained from each channel was evaluated via binomial test with the expected success probability set as 25%. Because multiple comparisons were done for each subject, the p-values obtained were FDR-adjusted ^63^.

### LFP analysis

Spectral content of the LFP data from cortical contacts of all six subjects and from hippocampal depth electrode recordings from three subjects was evaluated using EEGLAB (RRID:SCR_007292) routines that applied wavelet transforms ^64^ Spectral power was calculated over 1 to 200 Hz across each individual time period (“trial”) centered around ECoG HG peaks found in SWS patterns by convoluting each trial’s signal with complex Morlet wavelets, and averages were then taken across trials. The resulting time-frequency matrices were normalized with respect to the mean power at each chosen frequency and masked with two-tailed bootstrap significance, with the pre-trigger times (−1500 ms to −1000 ms) as baseline.

For automatic theta oscillation detection, data from SWS periods were bandpassed from 5-9 Hz, and the amplitude envelope from Hilbert transform was used alongside detection thresholds based on standard-deviation-over-mean ^36^. The theta events thus selected were further pruned by counting the number of zero-crossings over the duration of each event and calculating the zero-crossing-based frequency estimate, whereby if a given event’s estimate fell outside the bandpass range, it was removed from analysis. Automatic spindle detection was performed using previously published methods for SEEG spindle identification, with parameters tuned for cortical spindles ^65^: a 10-16Hz bandpass was applied to the data and the analytic amplitude of the band passed signal was convolved with an average Tukey window of 600 ms. A channel-wise cutoff set at mean +2 s.d. was applied to each channel’s convolved bandpass envelope to identify local maxima that would be “peaks” of putative spindles. To define the onset and offset of putative spindles, a 400 ms Tukey window was applied to the previously computed Hilbert amplitude, and the maximum FFT amplitude of the putative spindle signal at the spindle peak was used to set edge thresholds (50% of peak amplitude). Finally, the putative spindle segments were evaluated with four different criteria for rejection: 1, the spindle duration needs to be longer than 400 ms; 2, the spectral power outside the spindle frequency band on the lower end (4-8 Hz) or the higher end (18-30 Hz) should not exceed 14dB; 3, each spindle should consist of at least 3 unique oscillation peaks in the 10-16 Hz bandpass; 4, the time between successive zero-crossings should be within the range of 40-100 ms.

Hippocampal sharp-wave ripples were detected following earlier published methods ^37^, with some modifications. First, data from SWS were filtered between 80-100 Hz (6th order Butterworth IIR bandpass filter). Root-mean-square (RMS) over time of the filtered signal was calculated using a moving average of 20 ms, with the 99% percentile of RMS values for each hippocampal channel being set as a heuristic cut-off. Whenever a channel’s signal exceeded this cut-off continuously for at least 38 ms, a ripple event was detected. Since RMS peaks could arise from artifacts or epileptiform activity, 2000 milliseconds of hippocampal LFP data centered on each ripple event underwent 1-D wavelet (Haar and Daubechies 1-5) decomposition for the detection and removal of sharp transients (i.e. signal discontinuities). For each wavelet decomposition type, a scale threshold was established via iterative optimization for the best separation between ∼550 hand-marked true ripple events and ∼350 interictal events in the same SWS period. Automatic selections for theta bursts, spindles and ripples were visually reviewed for accuracy in the raw traces and confirmed for each subject and SWS period.

For our analyses involving graphoelements and Motifs/Events, we used Motif-Event matches that were obtained under the exact match (without shuffle) setup. To statistically evaluate the theta/spindle likelihood around all Motif-Event match HG peak times, we first made peristimulus histograms of theta/spindle occurrences with HG peaks as triggers (only peaks from Events that matched to waking Motifs and actually were part of the match were used). We used the midpoint of the theta/spindle occurrences as their times for histogram construction. For each HG peak, the theta/spindle counts to be added to the histogram only included the theta/spindle occurrences from the same channel. To obtain average histograms across multiple subjects, we first normalized individual subject’s histogram, dividing the count of each histogram bin by the total number of theta/spindle occurrences within the histogram time window (+-2s) to yield the observed likelihood for each subject, and averaged the observed likelihood across subjects. We then obtained the estimated average rates by multiplying the bin-wise average likelihood with the average number of total theta/spindle occurrences. Finally, we converted the counts into occurrence rates (dividing counts by the number of trials). We calculated the expected histograms under the null hypothesis that there is no relationship between Motif peak and theta/spindle occurrences: null hypotheses were constructed for consecutive time bins across the entire histogram (200 ms bins with 100 ms overlap): over each given time bin preceding/at/following Motif-Event match peaks. We then compared the observed likelihoods for each subject/graphoelement type to the numbers derived from 1000 sets of randomly initiated pseudo-HG peak locations in the same Sleep-Post, where the total number of HG peaks and the number of HG peaks from each channel remained the same as in the original, and the inter-peak intervals from real Motif-Event matches were preserved. The threshold for bootstrapping significance was set at 0.05, and the p-values thus obtained were FDR-adjusted.

Similarly, in order to statistically evaluate the ripple event likelihood around Motif-Event match HG peak times, we made peri-stimulus histograms of ripple occurrences with HG peaks as triggers, but using only the first peaks from Motif-Event matches and the histogram time window was expanded to +-40 s. The null hypothesis for ripple histograms, in turn, were constructed using 4 s bins with 50% overlap, and 1000 sets of random pseudo-HG peak locations, each set containing the same number of pseudo-HG peaks as the actual number of Motif-Event matches for the given Sleep-Post period.

## Acknowledgements

The authors would like to thank the following for their support: Qianqian Deng, Darlene Evardone, Chris Gonzalez, Don Hagler, Milan Halgren, Erik Kaestner, Adam Niese, Burke Rosen, Rachel Mak-McCully, and Anna Sargsyan. This work was supported by the U.S. Office of Naval Research (N00014-13-1-0672).

## Author Contributions

W. D. and E. E. performed surgical implantations; D. F., P. D., O. D., S. C. and T. T. recruited subjects, and collected data; I. S., S.C. and T.T. were involved in study design; E. H. and X. J. designed the study; X. J. and I. S. analyzed the data; X. J. and E. H. wrote the paper. All authors reviewed and approved the manuscript.

## Data Availability

Because clinical data sets are used in this study, raw electrophysiological recordings and neuroimaging files will be shared on a case-by-case basis in accordance with IRB agreements. All computer programs/scripts used in this study will be available upon request.

